# N6-methyladenosine demethylase FTO serves as an indicator for predicting prognosis and immunotherapy response in individuals with gastric cancer

**DOI:** 10.1101/2023.05.15.540747

**Authors:** Shiheng Jia, Heng Zhou, Lanxin Cao, Cheng Sun, Xue Yu, Yanshu Li, Kai Li

## Abstract

**Background:** N6-methyladenosine (m^6^A) RNA methylation is the most common chemical decoration in mammalian RNAs which exerts vital effects on numerous cellular processes. Recently, m^6^A regulators have been validated to participate in promoting immune evasion and act as prognostic factors in various cancers. Nevertheless, the predictive abilities of m^6^A regulators for the prognosis and immunotherapy response in gastric cancer (GC) remain indistinct.

**Methods:** Herein, The Cancer Genome Atlas (TCGA), Genotype-Tissue Expression (GTEx) database, The Human Protein Atlas (HPA), and a clinical GC cohort were applied for differential expression analysis, correlation analysis, survival analysis, and hazard model construction. Consensus clustering analysis was performed to authenticate the PD-L1 (CD274) expression, stemness features, immune cell infiltration, and tumor microenvironment (TME) in GC individuals. Furthermore, protein-protein interaction, immunotherapy response prediction, and drug susceptibility prediction were performed, respectively. Additionally, tissue microarray (TMA), immunohistochemical staining, western blot assay, Transwell assay, and flow cytometry assay were adopted to evaluate the protein expression, the prognostic value, and the influence of FTO on GC malignant phenotypes as well as the expression of PD-L1.

**Results:** In agreement with the majority of m^6^A regulators, FTO was overexpressed and predicted poor prognosis in GC. Based on consensus clustering analysis, two independent subgroups (G1/G2) were identified. Notably, FTO was upregulated in the G1 subgroup. Meanwhile, the infiltration level of CD8+ T cells was strikingly decreased while the stemness features were enhanced in the G1 subgroup. More importantly, FTO was negatively correlated with microsatellite instability (MSI) and tumor mutation burden (TMB). Furthermore, immune checkpoint blockade (ICB) response prediction indicated that patients with upregulated FTO showed high tumor immune dysfunction and exclusion (TIDE) scores. Subsequently, FTO was confirmed to be related to multiple immune checkpoints, particularly PD-L1. Specifically, FTO was dramatically upregulated in GC cell lines and clinical cancer samples. Functional experiments illustrated that FTO acted as an oncogene to facilitate malignant phenotypes. Notably, PD-L1 was remarkably downregulated after RNA interference-mediated knockdown of FTO.

**Conclusion:** FTO can aggravate GC malignant phenotypes. More importantly, it could be utilized to predict the long-term prognosis and the immunotherapy response in GC individuals. However, larger trials should be performed to verify the prediction accuracy.

## Introduction

Gastric cancer (GC) ranks as the fifth malignancy in tumor incidence and the fourth in cancer-related mortality worldwide[1]. Although the therapeutic strategies have been improved in GC patients, the prognosis remains poor due to tumor recurrence, distant metastasis, and therapeutic resistance. Notably, N6-methyladenosine (m^6^A) modification acts as a regulatory role in maintaining cancer hallmarks, particularly immune evasion[2]. Nowadays, m^6^A regulators have been noticed as hopeful molecular targets for cancer therapy[3]. Historically, m^6^A modification induced by methylation of adenosine at the N6 position was first detected in 1974[4]. Biologically, m^6^A methylation is the most abundant form among more than 170 internal RNA modifications identified at the post-transcriptional level[3] and participates in numerous physiological and pathological processes, including the initiation and aggressiveness of cancer[5–7]. Topologically, the dynamic reversible m^6^A methylation primarily presents in the consensus motif of RRACH (A = m^6^A; R = G or A; H = A, C, or U) which is localized in the vicinity of stop codons and 3′ untranslated regions (3′-UTRs)[8]. Functionally, m^6^A modification exerts vital effects on administrating the metabolism of RNA, involving RNA stability, translation, splicing, export, and decay[9]. Mechanistically, the canonical process of m^6^A methylation is dynamic and reversible. The initiation, recognition, and removal of m^6^A methylation are supervised by 3 types of proteins: m^6^A methyltransferases (“writers”), m^6^A binding proteins (“readers”), and m^6^A demethylases (“erasers”), respectively[10]. Specifically, m^6^A methylation is decorated by m^6^A writers, mainly involving METTL3, METTL14, and WTAP. METTL3 and METTL14 establish a multicomponent m^6^A methyltransferase complex (MTC) to install the m^6^A modification[11]. METTL3 binds to the S-adenosyl methionine (SAM) and catalyzes the transfer of methyl group by the hyperactive methyltransferase domain. METTL14 constructing a heterodimer with METTL3 plays a crucial role in m^6^A deposition and substrate recognition, facilitating the methylase activity[12]. Meanwhile, auxiliary proteins involving WTAP, VIRMA, ZC3H13, and RAB15/15B stabilize the MTC assembly. On the contrary, m^6^A methylation is demolished by m^6^A erasers, including FTO[13] and ALKBH5[14]. Indeed, m^6^A readers are real performers in the m^6^A process. Generally, m^6^A readers are mainly classified into three types, involving the YT521-B homology (YTH) domain family, heterogeneous nuclear ribonucleoproteins (HNRNPs), and IGF2BPs. Thereinto, FTO as an m^6^A demethylase can be utilized by tumors to aggravate malignant phenotypes and immune evasion. For instance, FTO-mediated m^6^A modification in tumor cells promotes metabolic reprogramming to escape immune surveillance[15, 16].

Recently, emerging evidence has demonstrated that immune evasion acts as a vital role in tumor survival and progression[17]. In general, tumor cells can establish a suppressive microenvironment by recruiting immunosuppressive cells to evade CD8+ T cell-mediated immune surveillance[18]. Strikingly, programmed death ligand 1 (PD-L1) interacting with its receptor programmed death-1 (PD-1) can enhance immune evasion by mediating tumor-specific T cells and inhibiting the activity of CD8+ T cells in cancers. Clinically, immunotherapy targeting immune checkpoint PD-1 and its ligand (PD-L1) has obtained extraordinary achievement in various cancers. However, the resistance of immune checkpoint inhibitors limits the persistence of immunotherapy. Additionally, the potency of PD-L1 expression in predicting anti-PD-L1 efficacy remains controversial. The KENOTE-061 and KENOTE-062 trials exhibited that pembrolizumab treatment can prolong the survival of individuals with PD-L1-positive tumors[19, 20]. However, the Checkmate032, JAVELIN Gastric 300, and ATTRCTION-2 trials showed an unsupportive viewpoint that PD-L1 positivity functioned as a predictive biomarker of anti-PD-1 efficacy[21–23]. Hence, the underlying mechanism of immune evasion in GC cells by monitoring PD-L1 expression needs further elaboration.

Herein, a comprehensive analysis was applied to predict the anti-PD-L1 efficacy, drug susceptibility, and prognosis in GC based on the expression of 20 m^6^A regulators. We clustered the GC samples in TCGA into two subtypes relying on the levels of m^6^A regulators and investigated the immune cell infiltration, cancer stemness, chemotherapeutic drug sensitivity, and the association between m^6^A regulators and immune checkpoints in the clustering subtypes. Additionally, we verified that silencing FTO can downregulate the expression of PD-L1, thereby promoting invasion, accelerating migration, and inhibiting apoptosis via in vitro assays. In general, our results provide novel perceptions for the prediction of immunotherapy response of PD-L1 in GC and identify a new target and therapeutic implication for GC treatment.

## Materials and Methods

### Data collection

Clinical information and RNA transcriptomes were obtained from the Genotype-Tissue Expression (GTEx) database and the Tumor Genome Atlas (TCGA) database. The databases included 375 GC tissues, 32 adjacent GC tissues, and 359 GTEx normal gastric epithelial tissues for further analysis. Clinical information was composed of gender, age, survival state, survival time, and Tumor Node Metastasis (TNM) stage.

### Exploration of canonical m^6^A regulators

In our study, 20 canonical m^6^A regulators were chosen for further research, involving METTL3, METTL14, YTHDF1, YTHDF2, YTHDF3, WTAP, VIRMA, RBM15, RBM15B, ZC3H13, FTO, ALKBH5, RBMX, IGF2BP1, IGF2BP2, IGF2BP3, YTHDC1, YTHDC2, HNRNPC, and HNRNPA2B1. Subsequently, the differential expression analysis of the aforementioned m^6^A regulators between GC and the control group was conducted and visualized by ggplot2 and pheatmap R packages. Moreover, Spearman’s correlation analysis was exploited to clarify the correlation between m^6^A regulators and displayed by the pheatmap R package. In addition, the protein-protein interaction (PPI) network of m^6^A regulators was established via the STRING database.

### Prognostic model construction and evaluation

Univariable and multivariable Cox hazard regression models constructed by the forestplot R package were utilized to estimate hazard ratios (HRs). And nomograms were built to predict 1, 3, and 5-year overall survival (OS) by rms R package. The least absolute shrinkage and selection operator (LASSO) regression was performed to obtain a set of prognostic m^6^A regulators and their LASSO regression coefficients by the glmnet R package. The risk score equation was the sum of coefficients * m^6^A regulator expression levels. Consequently, the risk score of individuals in TCGA-STAD was acquired. The median risk score was applied as the cutoff criterion to categorize the individuals into high-risk and low-risk groups. Kaplan–Meier analysis was utilized to evaluate the difference in OS between the high and low-risk groups by survival R package. Receiver operating characteristic (ROC) curves were performed to assess the capacity of the risk scores, and the area under the curve (AUC) was obtained.

### Consensus clustering analysis and signatures in clustering subgroups

The consensus clustering analysis was implemented using the ConsensusClusterPlus R package. The parameters were as follows: the best clustering stability from 2 to 6, clusterAlg = “hc”, innerLinkage = “ward.D2”. The clustering heat map was established by the pheatmap R package. CIBERSORT which detected the relative proportion of infiltrating immune cells in GC samples was utilized to assess the immune infiltration of clustering subgroups. The Genomics of Cancer Drug Sensitivity (GDSC) database was applied to predict the anti-tumor drug response by the pRRophetic R package. The half-maximal inhibitory concentration (IC50) was estimated by ridge regression and all parameters were set by default. The OCLR algorithm was adopted to calculate the mRNA stemness index (mRNAsi) to evaluate the tumor stemness in different subgroups.

### Differential expression analysis and immunotherapy response prediction

Differential expression analysis was accomplished by limma R package using level 3 RNA-sequencing data and corresponding clinical information in TCGA and GTEx databases and validated by The Human Protein Atlas at the protein level. Correlation analysis between FTO gene expression and TMB/MSI was performed using Spearman’s method. And potential ICB response was predicted via the TIDE algorithm. Correlation analysis between FTO and each immune checkpoint including SIGLEC15, TIGIT, CD274, HAVCR2, PDCD1, CTLA4, LAG3, and PDCD1LG2 was fulfilled based on mRNA expression levels.

### Tissue specimen collection

In this study, a clinical GC cohort of 287 GC patient tissues and 53 adjacent tumor controls was obtained from March 2007 to December 2008 at the Department of Surgical Oncology and General Surgery, the First Hospital of China Medical University. All aspects of this study were approved by the Medical Ethics Committee of the First Hospital of China Medical University, and all participating patients signed written informed consent. All collected tissues were immediately snap-frozen in liquid nitrogen and stored at -80°C.

### Cell lines and cell culture

Gastric epithelial cell lines (GES-1) and GC cells (HGC-27, MGC-803, MNK-45, and SNU-719) were kindly provided by Stem Cell Bank, Chinese Academy of Sciences. Cells were cultured using Dulbecco Modified Eagle medium (DMEM) (Gibco, Rockville, MD, USA) and RPMI 1640 (Biosharp, China). It contained 10% fetal bovine serum (FBS) (VivaCell Biosciences, Shanghai, China) and was incubated at 37℃ in a 5% CO2 incubator. Cells were passaged using trypsin until the cells grew to 70-80% confluence (BI, Waltham, MA, USA).

### Cell transfection

Transfection RNA oligo was purchased from GenePharma (Shanghai, China). Cells were cultured to 30-50% confluence and transfected using HieffTrans TM Liposomal Transfection Reagent (Shanghai, China). Transfected cells were assembled 72h later for subsequent use. All of the siRNA nucleotide sequences were listed in table 2 in supplementary file .

### Transwell assay

250 μL of the suspension (2 × 10 ^5^ cells/mL) was applied to the upper layer of a Transwell chamber (Millipore, Billerica, MA, USA), which was placed in a 24-well plate containing 800 μL of 10% FBS at the bottom. After 24 hours of incubation, the bottom cells were reacted with methanol for 20 minutes, then crystal violet for 30 minutes, and subsequently captured by microscopy. Count migrating cells for each sample in 5 random fields.

### Western blot assay

Cells were lysed to separate cellular proteins and run for electrophoresis. Protein samples were transferred onto polyvinylidene fluoride (PVDF) membranes (Millipore, Billerica, MA, USA). After, non-specific antigens were blocked in 3% BCA for 1 hour. The membrane was reacted with the primary antibody at 4℃ overnight. After recovering to room temperature for 30 minutes, TBST was used to wash three times for five minutes each and then reacted with the secondary antibody for one hour at ordinary temperature. FTO (1:1000; ab12660; Abcam), PDL1 (1:1000; ab205921; Abcam), GAPDH (1:1000; ab8245; Abcam) were purchased and used in this study. Finally, band exposure and analysis were performed.

### RNA Extraction and Real-Time PCR Assay

First, TRIzol (Invitrogen, Carlsbad, CA, USA), chloroform, and isopropanol were used to extract total RNA from cell lines and gastric tissues. Then, according to the instructions, a reverse transcription kit (TaKaRa, Shiga, Japan) was applied to reverse transcribe 1000ng total RNA into cDNA. SYBR dyes (Takara, Shiga, Japan) were combined into a 12ul system for cDNA amplification and signal collection. Using the Mx3000PTM real-time PCR system of Agilent (Stratagene, La Jolla, CA, USA), mRNA levels of FTO and internal reference GAPDH were measured by real-time quantitative PCR in triplicate.The specific primers used for these genes are listed in table 1 in supplementary file.

### Flow cytometry assay

Harvested GC cells and resuspended in 495 μL of 1× binding buffer. First, added 5 μL Annexin V-PE to the cell suspension, then added 10 μL 7-AAD (Annexin V-PE/7-AAD Apoptosis Detection Kit, KeyGEN Biotechnology, Shanghai, China). Finally, cells were mixed well and incubated in the refrigerator at 4°C for 15 minutes in the dark. Cells were detected by flow cytometry (Accuri C6, Becton-Dickinson, US).

### TMAs and IHC methods

All GC tissue microarrays were cut into 3 μm sections, dewaxed and hydrated, and then used with UltraSensitivity TM S-P system (KIT-9720; Mai X, Fuzhou, China) and were incubated for 24 hours at 4°C with anti-FTO antibody (1:200, #sc-271713; SANTA, Cambridge, MA, USA). After washing with PBS three times for 8 minutes each time, a biotin-labeled secondary antibody was used to react with tissue sections for 30 minutes at room temperature. Tetrahydrochloride diaminobenzidine substrate (Maixin) was used as the chromotropic reagent, and the chromotropic reagent was washed with tap water for one minute immediately after successful chromotropic development. The FTO expression and staining scores were as follows: 0 (no staining), 1 (weak expression), 2 (moderate expression), and 3 (high expression). The dyeing percentage score distribution was as follows: 0 (< 10%), 1 (10%-25%), 2 (26%-50%), 3 (51% -75%), and 4 (76% -100%). The score of each tumor sample was multiplied to gain a final score of 0-12, and the staining results were classified as negative (0; −), low (1-4; +), medium (5-8; + +), and high (9-12; + + +). Each patient’s tissue sections were viewed in 10 fields, and sections were read by two experienced pathologists without knowledge of clinical information and pathological grade. The experimental results were determined by the double-blind method.

### Statistical Analysis

Statistical analysis was performed using R software (v4.0.3). The Student’s t-test, Wilcoxon test, and Kruskal-Wallis test were used to determine differences between two groups and among several groups, respectively. Spearman’s correlation analysis was performed to assess the subgroups, clinical features, risk score, expression of FTO and PD-L1, and immune infiltration levels. Kaplan–Meier analysis and a log-rank test were utilized to distinguish the OS between the two groups. P < 0.05 was reckoned statistically significant.

## Results

### The m^6^A regulators with synergistic effects are differently expressed in GC

The differential expression of 20 m^6^A regulators was initially outlined depending on the transcriptome data in TCGA and GTEx. METTL3, METTL14, WTAP, VIRMA, RBM15, RBM15B, ZC3H13, FTO, ALKBH5, RBMX, IGF2BP1, IGF2BP2, IGF2BP3, YTHDC1, YTHDC2, YTHDF1, YTHDF2, YTHDF3, HNRNPC, and HNRNPA2B1 were significantly overexpressed in GC individuals compared to normal control (Figure 1A, 1B). Then, we analyzed the correlation of these m^6^A regulators in the TCGA-STAD samples. Among the m^6^A writers, METTL3 was significantly and positively correlated with RBMX, YTHDC1, YTHDC2, YTHDF2, YTHDF3, HNRNPC, and HNRNPA2B1. METTL14 was positively correlated with YTHDC1, YTHDC2, and YTHDF3. WTAP was positively correlated with RBMX, YTHDC1, YTHDC2, and YTHDF3. VIRMA and RBM15 were positively correlated with RBMX, YTHDC1, YTHDF1, YTHDF2, YTHDF3, and HNRNPA2B1. ZC3H13 was positively correlated with YTHDC1 and HNRNPA2B1. Both erasers including FTO and ALKBH5 were positively correlated with YTHDC1 (Figure 1C). The PPI network of 20 m^6^A regulators indicated that m^6^A writers, readers, and erasers can interact with each other, which verified the dynamic and reversible m^6^A process (Figure 1D).

**Figure 1.**
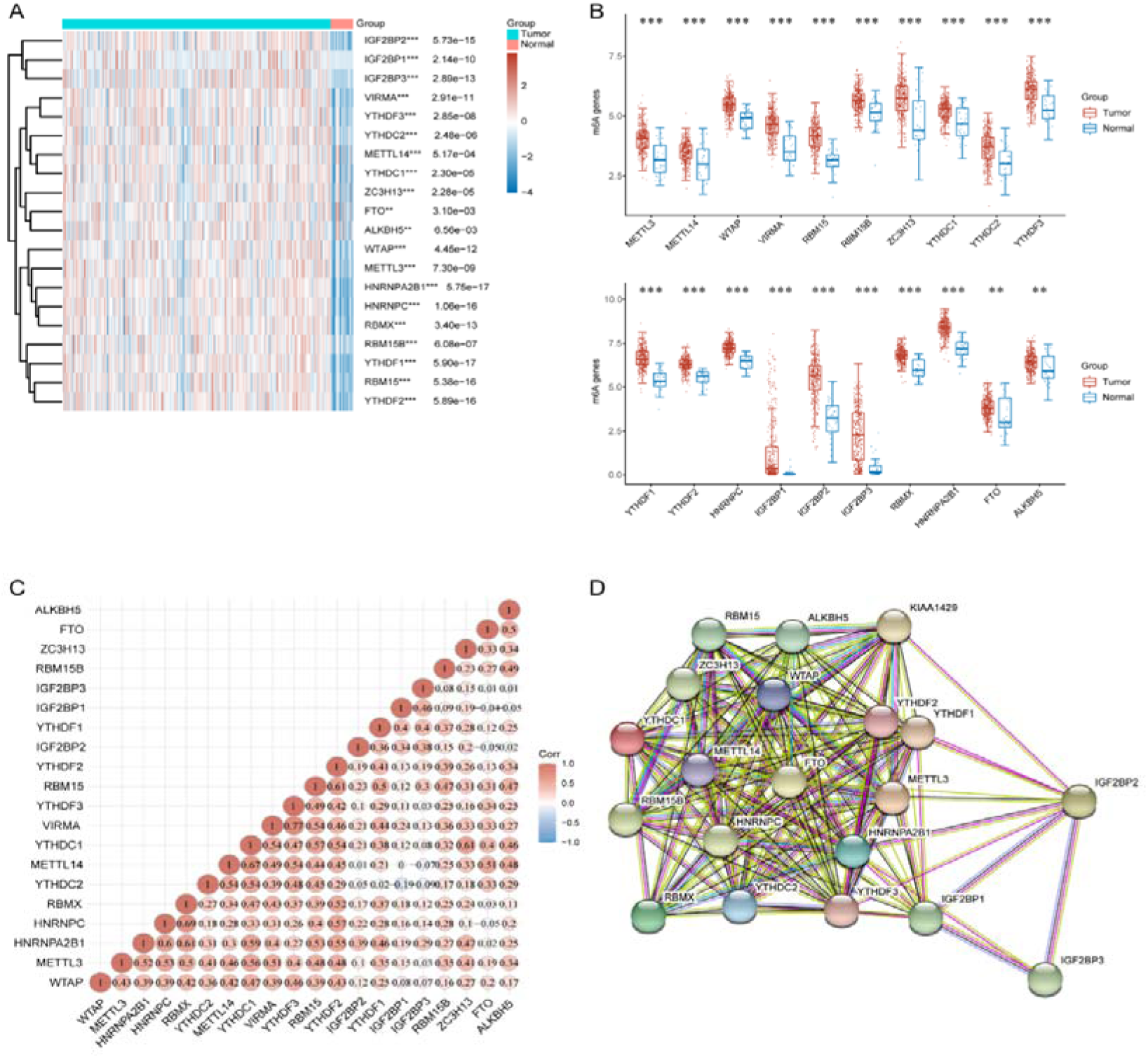
Expression, interaction, and correlation of m^6^A regulators in GC. **(A)** Heat map illustrating the dysregulated m^6^A regulators in GC samples. (** P < 0.01, *** P < 0.001). **(B)** Histogram illustrating differential expression of m^6^A regulators in GC and adjacent tissues. All m^6^A RNA methylation regulators were significantly overexpressed in GC individuals compared to normal control (** P < 0.01, *** P < 0.001). **(C)** The correlation of m^6^A regulators in the TCGA-STAD samples was analyzed by Pearson’s correlation. Both erasers including FTO and ALKBH5 were positively correlated with YTHDC1. **(D)** The PPI network of 20 m^6^A regulators indicated that m^6^A writers, readers, and erasers can interact with each other.

### A prognostic model is established to predict the survival of GC patients

Univariate and multivariate cox regression analyses were utilized to ascertain the appropriate terms to construct the nomogram (Figure 2A, 2B). Univariate Cox regression analysis demonstrated that FTO (p = 0.027, HR = 1.384), RBM15 (p = 0.011, HR = 0.677), Age (p = 0.009, HR =1.021), and pTNM stage (p = 0.008, HR = 1.274) were significantly associated with OS (Figure 2A). Subsequently, multivariate Cox regression analysis illustrated that FTO (p = 0.004, HR = 1.693), RBM15 (p = 0.002, HR = 0.511), Age (p = 0.0001, HR =1.034), and pTNM stage (p = 0.021, HR = 1.280) were independent prognostic factors for GC (Figure 2B). Nomograms to predict 1-, 3-, and 5-year OS were constructed. The C-index (Figure 2C) and calibration plots (Figure 2D) presented a good discriminative ability and an optimal accuracy of the nomograms. LASSO regression was utilized to calculate the coefficient of FTO and RBM15 (Figure 3B), and risk score = (0.4098 * FTO expression) + (-0.4823 * RBM15 expression), lambda. min=0.009. According to the risk scores, GC patients in TCGA were divided into two dichotomous groups (Figure 3A, 3B). Overall, the individuals in the low-risk score group displayed a longer OS than those in the high-risk score group (Figure 3C). Additionally, Time-dependent ROC curves were plotted to assess the predictive accuracy of this model, and The AUC of 1-, 3-, and 5-year OS was 0.62, 0.661, and 0.668, respectively (Figure 3D).

**Figure 2.**
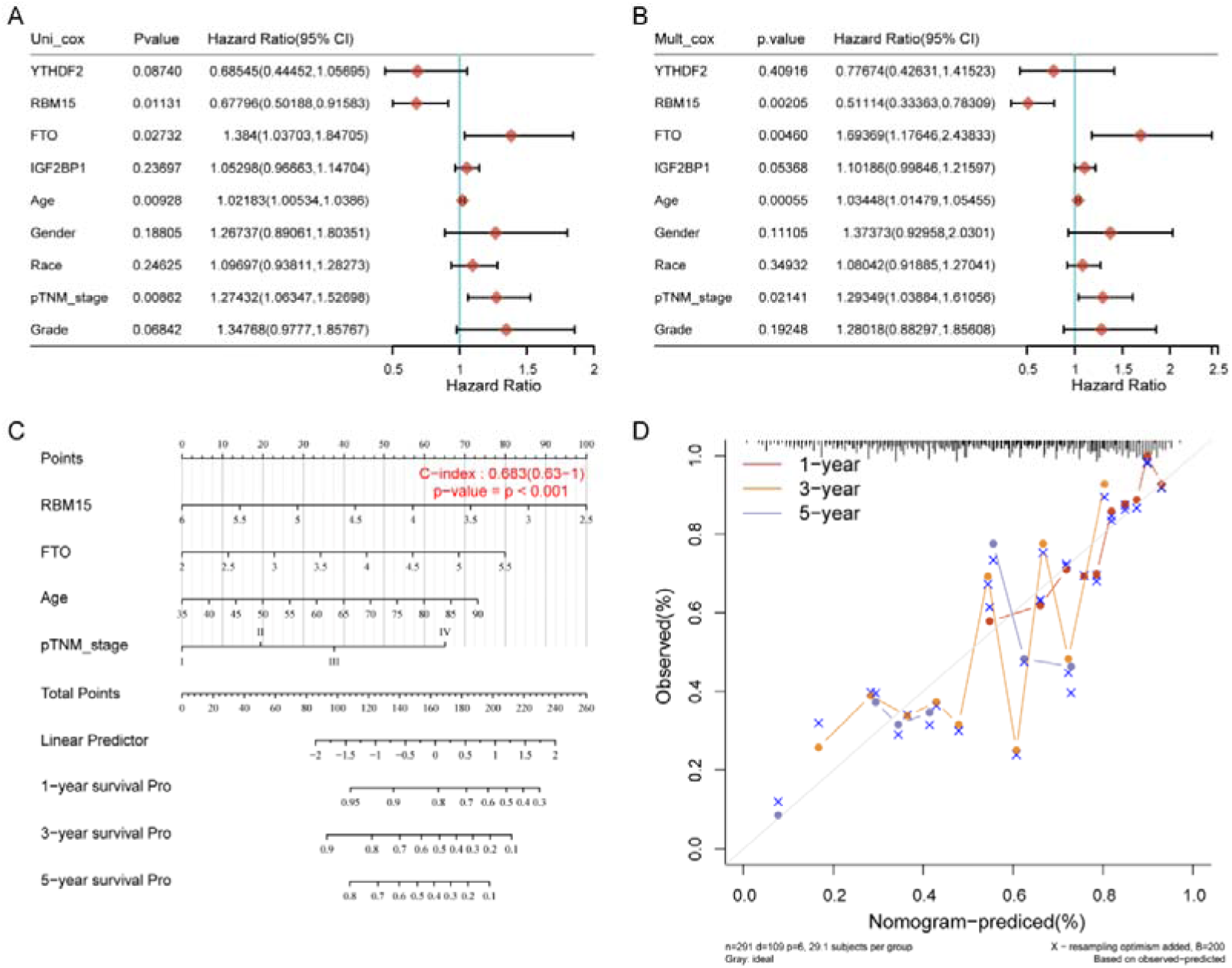
Construction of the prognostic signature based on the TCGA GC cohort. **(A, B)** Univariate and multivariate cox regression analyses were utilized to ascertain the appropriate terms to construct the nomogram. FTO and RBM15 were independent prognostic factors for GC and were selected to construct the nomogram. **(C, D)** Nomograms to predict 1-, 3-, and 5-year OS were constructed. The C-index and calibration plots presented a good discriminative ability and an optimal accuracy of the nomograms.

**Figure 3.**
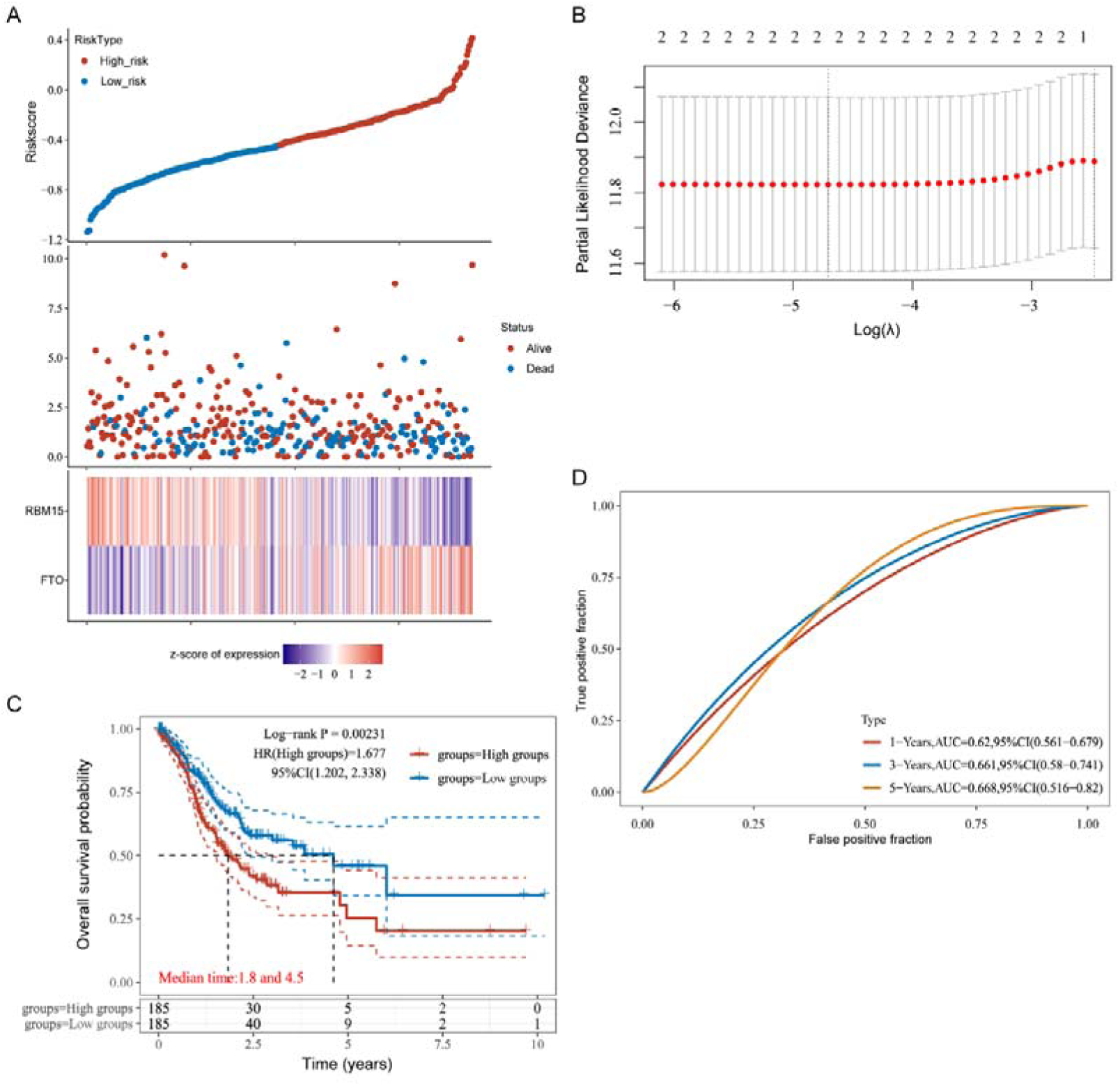
The relationship of prognostic model based on m^6^A regulators and OS in GC. **(A, B)** The prognostic signatures were constructed by the minimum criterion of the LASSO Cox regression algorithm. According to the risk scores, GC patients in TCGA were divided into two dichotomous groups. **(C)** Kaplan Meier curve showed that the prognostic model based on m^6^A regulators was significantly correlated with OS in GC. The individuals in the low-risk score group displayed a longer OS than those in the high-risk score group. **(D)** Time-dependent ROC curves were plotted to assess the predictive accuracy of this model, and the AUC of 1-, 3-, and 5-year OS was 0.62, 0.661, and 0.668, respectively.

### The m^6^A regulators exhibit different immune signatures, drug susceptibility, and stemness characteristics in the subgroup of GC

Based on the expression similarity of m^6^A regulators, k = 2 was demonstrated to be the most appropriate selection to divide the 375 GC patients into G1 and G2 subgroups (Figure 4A-4D). The results of the differential analysis indicated that all the m^6^A regulators except IGF2BP1 and IGF2BP3 were overexpressed in the G1 subgroup (Figure 4E).

**Figure 4.**
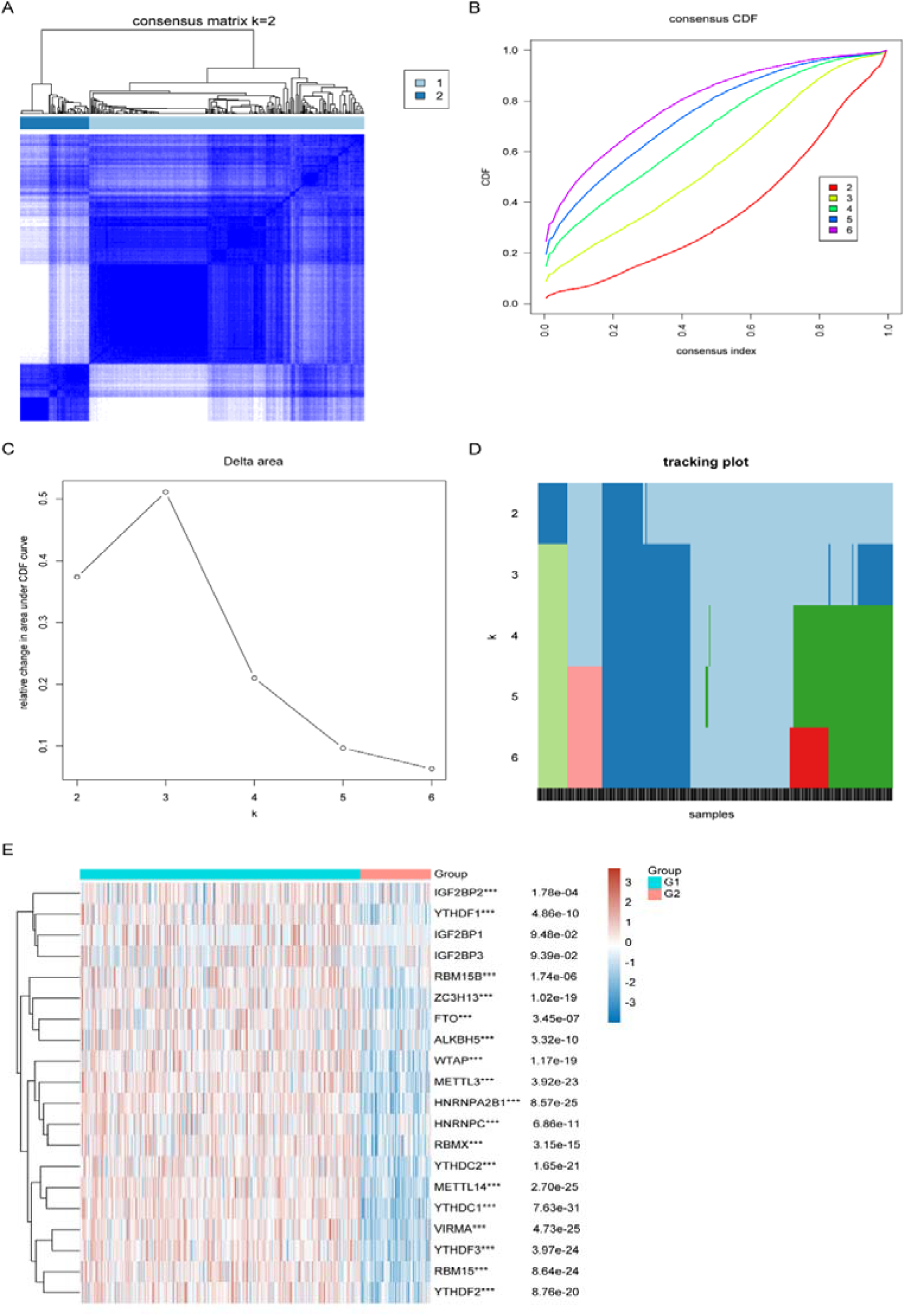
Two GC patient subgroups were identified by consensus clustering. **(A-D)** The GC cohort from TCGA was divided into two distinct clusters when k = 2. **(E)** The results of the differential analysis indicated that all the m^6^A regulators except IGF2BP1 and IGF2BP3 were overexpressed in the G1 subgroup (** P < 0.01, *** P < 0.001).

The analysis of immune cell infiltration revealed that the G1 subgroup showed higher levels of B cell naive, and T cell CD4+ memory, whereas the G2 subgroup displayed higher levels of B cell memory, B cell plasma, T cell CD8+, T cell regulatory (Tregs), and Monocyte (Figure5A, 5B). The prediction of drug susceptibility manifested that the patients in the G1 subgroup were sensitive to Docetaxel (Figure 5C) and Paclitaxel (Figure 5D), while the patients in the G2 subgroup were sensitive to Cisplatin (Figure 5E), Gemcitabine (Figure 5F), and Doxorubicin (Figure 5G). Additionally, the prediction of tumor stemness by mRNAsi suggested that the stemness in the G1 subgroup was significantly increased compared to the G2 subgroup (Figure. 5H).

**Figure 5.**
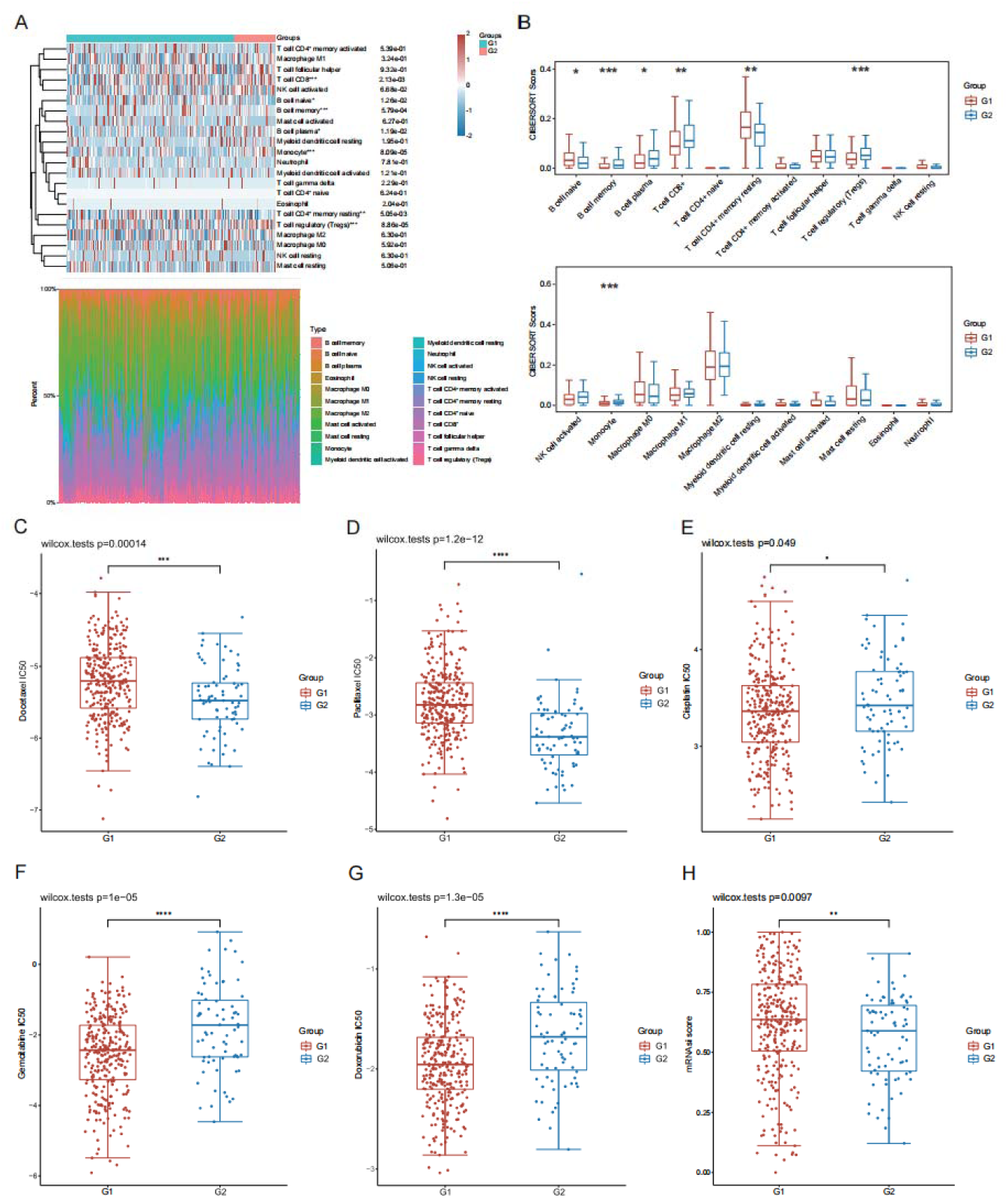
Comparison of the relationship between the immune signatures, drug susceptibility, and stemness characteristics in two clusters. **(A, B)** Comparison of the relationship between the immune signatures of two clusters. The G1 subgroup showed higher levels of B cell naive, and T cell CD4+ memory, whereas the G2 subgroup displayed higher levels of B cell memory, B cell plasma, T cell CD8+, T cell regulatory (Tregs), and Monocyte (* P < 0.05, ** P < 0.01, *** P < 0.001). **(C-G)** Comparison of the relationship between the drug susceptibility of two clusters. the patients in the G1 subgroup were sensitive to Docetaxel and Paclitaxel, while the patients in the G2 subgroup were sensitive to Cisplatin, Gemcitabine, and Doxorubicin (* P < 0.05, ** P < 0.01, *** P < 0.001, **** P < 0.0001). **(H)** The difference in stemness characteristics between the two groups (** P < 0.01).

### Overexpressed FTO is related to worse survival, MSI, TMB, ICB response, and upregulated PD-L1

FTO was upregulated in GC samples and acted as a crucial prognostic factor as described above. The overexpression of FTO protein was confirmed via The Human Protein Atlas (Figure 6A). Interestingly, FTO also presented higher expression in the T4 stage compared to T2 and T3 stages in GC (Figure 6B), indicating that FTO may promote cancer progression. Meanwhile, FTO was verified as an independent prognostic biomarker in GC patients by the Kaplan-Meier plotter database which was capable to evaluate the correlation between the expression of genes and survival in 30k+ samples from 21 tumor types (Figure 6C). The limma package of R software was used to detect the differential expression of mRNA. The adjusted P value was analyzed in TCGA or GTEx to correct false positive results. The gene expression was divided into a high expression group and a low expression group by a median method according to the expression value. Then we performed the correlation analysis between FTO and immune checkpoints in the TCGA and GTEx databases and confirmed that FTO was positively correlated with numerous immune checkpoints (Figure 6D), particularly PD-L1 (Figure 6E). Hence, we analyzed the differential expression of PD-L1 in the FTO upregulated and downregulated groups and found that the FTO upregulated group showed a higher level of PD-L1 (Figure 6F). All the above indicated that FTO as an oncogene might participate in governing the expression of PD-L1. Notably, we detected the association between FTO expression and TMB, MSI, and ICB response, which are considered predictors of the effectiveness of tumor immunotherapy, and validated that FTO was negatively correlated with the TMB, MSI, and ICB response (Figure 6G, 6H, 6I), which means that FTO may be a potential predictor for ICB response.

**Figure 6.**
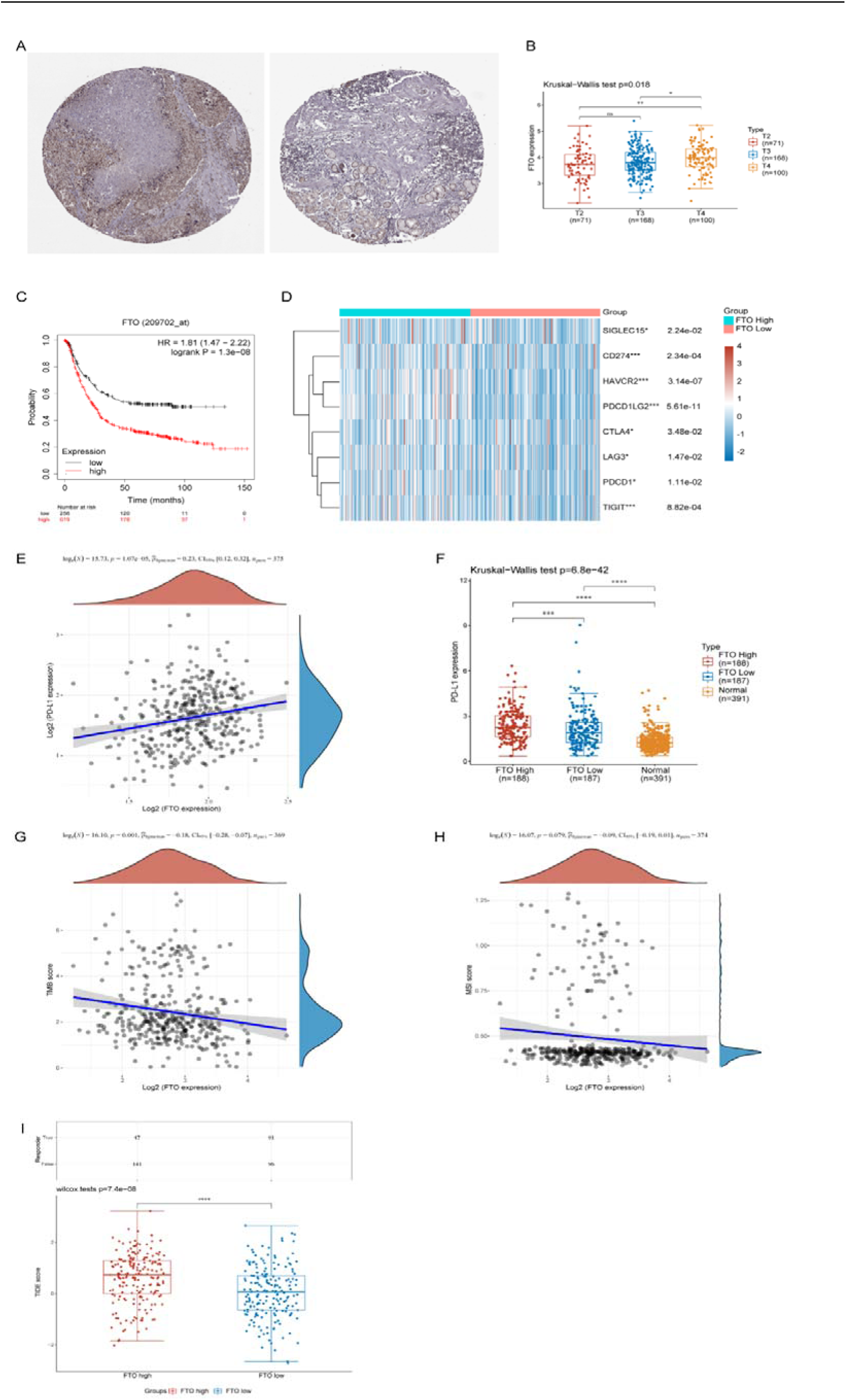
Overexpressed FTO is related to overall survival, MSI, TMB, ICB response, and upregulated PD-L1. **(A)** The overexpression of FTO protein was confirmed via The Human Protein Atlas. Compared with paracancerous tissue, FTO was over-expressed in cancer tissue. **(B)** FTO also presented higher expression in the T4 stage compared to T2 and T3 stages in GC (* P < 0.05, ** P < 0.01). **(C)** Kaplan-Meier plotter database was used to verify FTO as an independent prognostic biomarker in GC patients. **(D)** The gene expression was divided into a high expression group and a low expression group by the median method according to the expression value. The heat map showed that FTO was positively correlated with numerous immune checkpoints (* P < 0.05, *** P < 0.001). **(E)** FTO was significantly positively correlated with PD-L1. **(F)** GC patients with high expression of FTO in the TCGA database presented higher expression of PD-L1 (*** P < 0.001, **** P < 0.0001). **(G-I)** FTO was negatively correlated with the TMB, MSI, and ICB response (P =0.001, P =0.079, P < 0.0001).

### FTO is overexpressed and correlated with prognosis in the clinical GC cohort

To identify the dysregulated FTO in the tumorigenesis and progression of GC, we performed multiple analyses involving clinical GC cohort and GC cell lines. Tissue microarrays including 287 GC patients with well-characterized follow-up information were applied to evaluate FTO expression by immunohistochemistry. Notably, a remarkable overexpression of FTO was identified in GC samples (Figure 7A). Furthermore, Kaplan–Meier plot analysis was applied to evaluate the independent prognostic capacity of FTO and indicated that high expression of FTO was associated with shortened OS (Figure 7B). Subsequently, to evaluate the impact of each clinical variable on OS, we used univariate and multivariate Cox regression, as shown in Table 1. In univariate analysis, factors significantly associated with OS included tumor size (HR=0.343, 95% CI=0.160-0.732, P<0.001), age (HR=1.405, 95% CI=1.026-0.1.925, P<0.005), and FTO expression (HR=0.477, 95% CI=0.306-0.744, P=0.001). In multivariate analysis, the expression of FTO (P<0.01), and tumor size (P<0.01) were still associated with poor OS.

**Figure 7.**
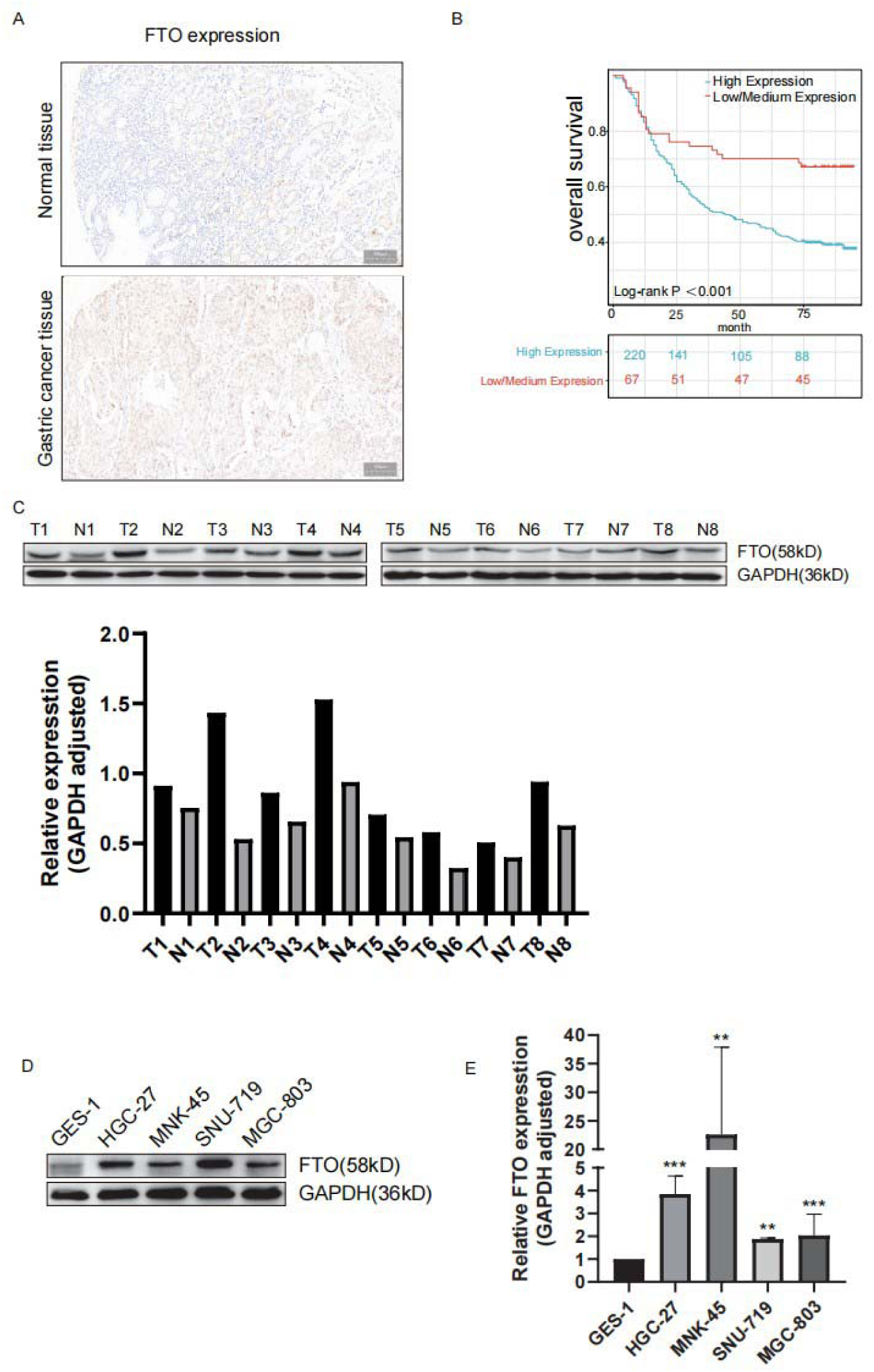
FTO is overexpressed and correlated with prognosis in the clinical GC cohort. **(A)** IHC was performed to measure the expression of FTO in GC tissues and adjacent normal tissues. Compared with paracancerous tissue, FTO was significantly over-expressed in cancer tissue. **(B)** Kaplan–Meier plot analysis was applied to evaluate the independent prognostic capacity of FTO. High expression of FTO was associated with shortened OS (P <0.001). **(C)** Western blot was utilized to detect the differential expression of FTO at the protein level in eight pairs of GC tissues and the corresponding paracancerous tissues. And FTO was highly expressed in the cancer tissues. **(D)** Western blot was performed to measure the expression of FTO in cell lines involving GES-1, HGC-27, MGC-803, MNK-45, and SNU-719. And FTO was upregulated in GC cell lines. **(E)** qPCR was performed to measure the expression of FTO in cell lines involving GES-1, HGC-27, MGC-803, MNK-45, and SNU-719, suggesting that FTO was upregulated in GC cell lines (** P < 0.01, *** P < 0.001).

**Table1.** Univariate and Multivariate Analysis of Overall Survival in GC

| Characteristics | Total(N) | Univariate analysis |  | Multivariate analysis |  |
| --- | --- | --- | --- | --- | --- |
|  |  | Hazard ratio (95% CI) | P value | Hazard ratio (95% CI) | P value |
| Age | 287 |  | **0.034** |  |  |
| <60 years | 152 | Reference |  | Reference |  |
| ≥60years | 135 | 1.405 (1.026 - 1.925) | **0.034** | 1.219 (0.881 - 1.687) | 0.232 |
| Sex | 287 |  | 0.461 |  |  |
| Male | 206 | Reference |  |  |  |
| Female | 81 | 0.877 (0.615 - 1.249) | 0.466 |  |  |
| Tumor size | 287 |  | **0.001** |  |  |
| >2cm | 260 | Reference |  | Reference |  |
| ≤2cm | 27 | 0.343 (0.160 - 0.732) | **0.006** | 0.327 (0.153 - 0.698) | **0.004** |
| FTO Expression | 287 |  | **< 0.001** |  |  |
| High Expression | 220 | Reference |  | Reference |  |
| Low/Medium Expression | 67 | 0.477 (0.306 - 0.744) | **0.001** | 0.497 (0.314 - 0.785) | **0.003** |

Moreover, we extracted eight pairs of GC tissues and the corresponding paracancerous tissues for western blot to detect the differential expression of FTO at the protein level. Consistently, FTO was highly expressed in the cancer tissues (Figure 7C). Additionally, the expression of FTO protein in cell lines involving GES-1, HGC-27, MGC-803, MNK-45, and SNU-719 was confirmed by western blot and q-PCR, suggesting that FTO was upregulated in GC cell lines (Figure 7D, 7E). Collectively, these findings demonstrated that FTO was dramatically overexpressed in GC and acted as a potential independent prognostic indicator for GC patients.

### The impact of FTO on biological characteristics of GC cells

To identify the biological functions of FTO participating in GC cell proliferation and progression, we initially performed Transwell assays to detect the influence of FTO on the invasive capacity of GC cells. Consequently, silencing FTO significantly suppressed the capacity for invasion and migration of HGC-27 and MGC-803 (Figure 8A). Furthermore, the depletion of FTO contributed to enhanced cell apoptosis of HGC-27 and MGC-803 via flow cytometry (Figure 8B). Taken together, these results suggested that FTO acts as an oncogene and maintains the oncogenic features in GC development and progression.

**Figure 8.**
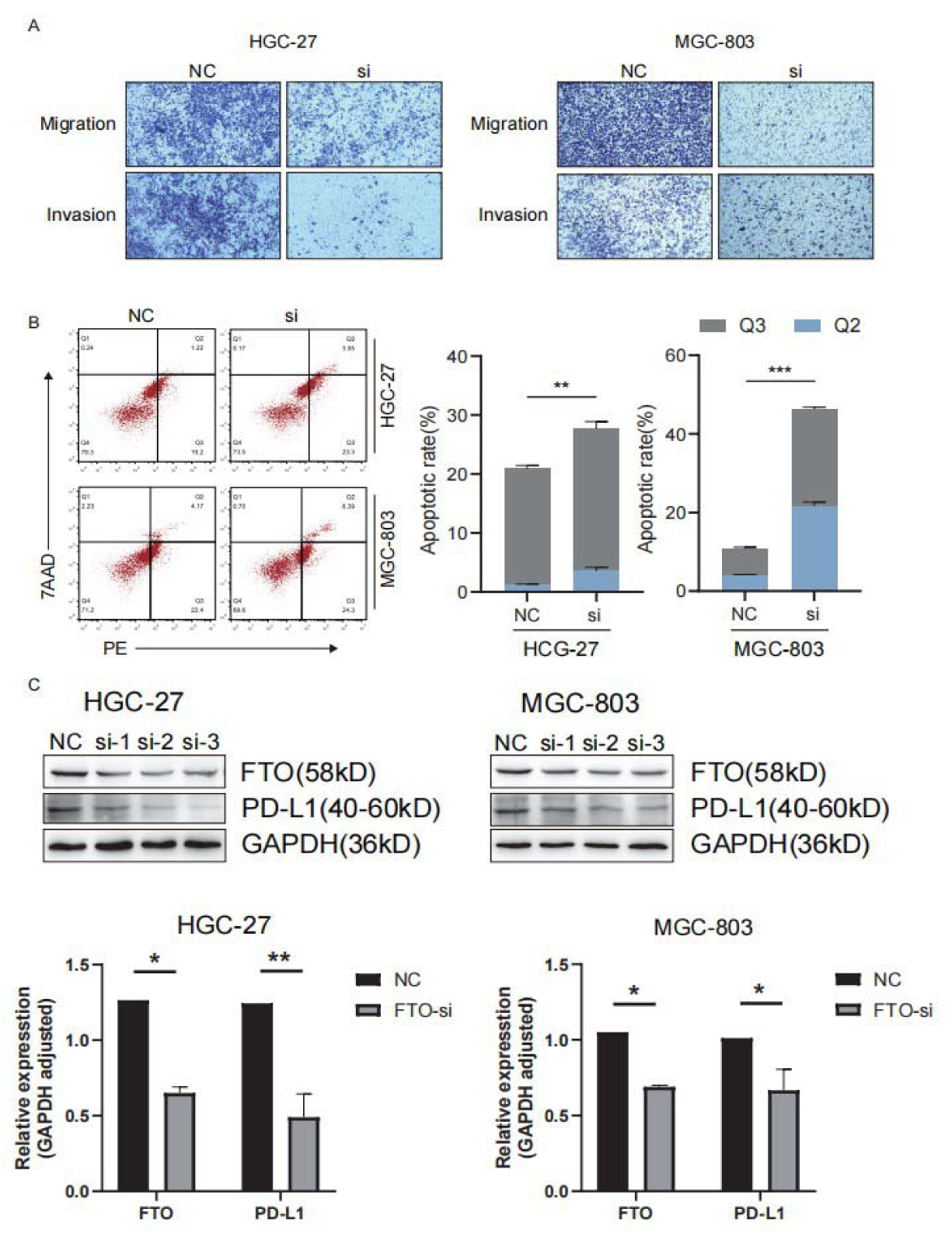
The impact of FTO on the biological characteristics of GC cells and the identification of the potential interaction between FTO and PD-L1 in GC cells. **(A)** Transwell assays to detect the influence of FTO on GC cell invasive capacity. Silencing FTO significantly suppressed the capacity for invasion and migration of HGC-27 and MGC-803. **(B)** Flow cytometry assay to detect the apoptosis of depletion of FTO in GC cell lines. The depletion of FTO contributed to enhanced cell apoptosis of HGC-27 and MGC-803 (** P < 0.01, *** P < 0.001). **(C)** Silencing FTO remarkably led to the decrease of PD-L1 in HGC-27 and MGC-803 cell lines (* P < 0.05, ** P < 0.01).

### Identification of the potential interaction between FTO and PD-L1 in GC cells

As described above, FTO was highly correlated with PD-L1 by certification in public databases. We speculated that FTO may involve in the regulation of PD-L1 expression. Therefore, we utilized the RNA interference assay to excavate the potential interaction between FTO and PD-L1. All 3 siRNAs presented satisfactory knock-down efficiency. As a result, the knockdown of FTO remarkably led to the decrease of PD-L1 in HGC-27 and MGC-803 cell lines (Figure 8D). Therefore, we validate the regulatory effect of FTO on the turnover of the PD-L1 protein.

## Discussion

GC is induced by numerous risk factors including environmental exposure, genetic mutations, and epigenetic modifications[6, 24, 25]. Strikingly, epigenetic modifications involving noncoding RNA regulation, histone modification, DNA methylation, and m^6^A modification perform etiologic functions to promote the pathological processes in GC[26]. Recently, emerging evidence has confirmed that m^6^A modification participates in the occurrence and development of multiple cancers[27]. Thereinto, FTO is frequently dysregulated and exerts an oncogenic role in maintaining cancer hallmarks in various cancers. For instance, elevated FTO promotes the growth and metastasis of clear cell renal cell carcinoma by mediating autophagy[28]. Moreover, FTO significantly facilitates the proliferation, migration, and invasion of multiple myeloma in an m^6^A-dependent manner[29]. Additionally, Upregulated FTO enhances the growth and metastasis of osteosarcoma via Wnt signaling. Intriguingly, FTO has been validated to maintain a strong correlation to the body mass index, obesity risk, and type 2 diabetes. Meanwhile, obesity and diabetes are risk factors for gastrointestinal tumors. Of note, a recent study has demonstrated that obesity can accelerate immune evasion of non-small cell lung carcinoma via upregulation of immune checkpoint Siglec-15 and glycolytic reprogramming[30]. Therefore, whether FTO is involved in the immune evasion of GC by targeting immune checkpoints needs further clarification.

PD-L1 is a significant immune checkpoint that participates in the inactivation of T cell proliferation and the inhibition of infiltration of CD8 and CD3 T cells into tumors [17]. PD-L1 inhibitor as an immune checkpoint blockade (ICB) exhibits promising benefits for cancer patients. But the high resistance rates limit its clinical application. Notably, m^6^A modification has been considered a vital regulator of the immune microenvironment and a considerable target for improving the efficacy of cancer immunotherapy. For example, a small-molecule ALKBH5 inhibitor, ALK-04, has been identified to enhance the efficacy of cancer immunotherapy[31]. In addition, two FTO inhibitors remarkably promote the immune response by restraining the expression of the immune checkpoint LILRB4 in leukemia[32]. Importantly, silencing the FTO sensitizes melanoma cells to interferon-gamma and anti-PD-1 therapy, which indicates that FTO facilitates the resistance of cancer immunotherapy[33].

Our present study confirmed that FTO was significantly overexpressed in TCGA-STAD and clinical GC cohort, which played a prognostic role in GC. Furthermore, this study constructed a prognostic risk model of GC from the perspective of m^6^A modification. The established prognostic risk model can predict the prognosis of GC individuals effectively. Meanwhile, consensus clustering analysis was utilized to divide GC individuals into two subgroups based on the expression signature of m^6^A-related genes. Notably, lower levels of T cell CD8+ and higher levels of tumor stemness were observed in the G1 subgroup in which most m^6^A regulators were upregulated. It is indicated that the G1 subgroup was more prone to immune evasion and chemotherapeutic resistance. This is similar to the previous research results of Ji H et al. in cervical cancer[33]. Subsequently, we confirmed the relationship between FTO expression and TMB, MSI, and ICB response. We validated that individuals with high expression of FTO showed higher TIDE scores and FTO expression was negatively correlated with TMB and MSI, which indicated that FTO may play a predictive factor for ICB response. We focused on the impact of overexpressed FTO on maintaining cancer hallmarks, including invasion, migration, apoptosis, and immune evasion. As expected, we identified that FTO facilitated invasion and migration, and inhibited apoptosis. Additionally, FTO was confirmed to be associated with multiple immune checkpoints in GC, especially PD-L1, which further suggested that FTO may be a potential target for immunotherapy.

In conclusion, our study systematically analyzed the expression profiles of m^6^A regulators in GC and their correlation with prognosis, immune infiltration, immune checkpoints, tumor stemness, and drug susceptibility, and constructed a prognostic risk model. FTO was confirmed to be a potential therapeutic target and a predictor of immune therapy in GC. Mechanistically, FTO promotes growth, invasion, and immune evasion by mediating the expression of PD-L1, which was validated by bioinformatics analysis and basic experiments. The application of PD-L1 inhibitors in GC still has many limitations. Whether GC patients can benefit from PD-L1 inhibitor treatment still has no precise indicator. The regulatory relationship between FTO and PD-L1 in GC, the correlation between FTO and TMB, MIS, ICB response validated by bioinformatics and basic experiments, suggest that FTO may become a novel potential predictor and target for the treatment of GC, but its regulatory mechanism still needs to be further explored.

## Data Availability Statement

The patients’ clinical data used to support the findings of this study are restricted by the The First Hospital of China Medical University in order to protect patient privacy. Data are available from the corresponding author upon request. The datasets analyzed in this study could be found in Genotype-Tissue Expression (GTEx) database https://commonfund.nih.gov/gtex and the Tumor Genome Atlas (TCGA) database.https://www.cancer.gov/ccg/research/genome-sequencing/tcga

## Ethics statement

The studies involving human participants were reviewed and approved by The Medical Ethics Committee of the First Hospital of China Medical University, and the written informed consent form to participate was signed by all the involved patients.

## Author Contributions

All authors contributed to the article and approved the submitted version.

## Supplementary

Table 1 shows the primer sequence and siRNA nucleotide sequence of FTO Table 2 shows the siRNA nucleotide sequence of FTO

---

**Table 1.**
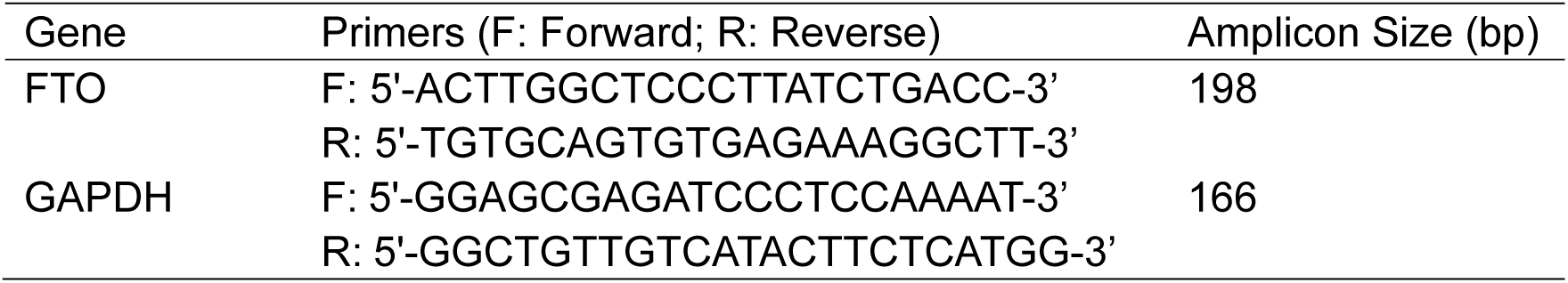
Sequence of Primers

**Table 2.**
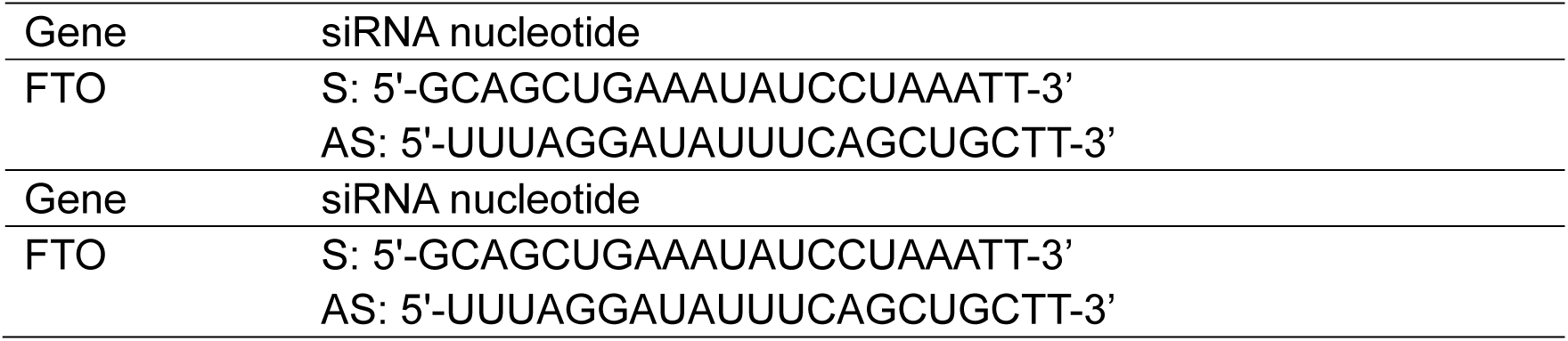
Sequence of siRNA nucleotide

